# Accelerated Molecular Dynamics and AlphaFold Uncover a Missing Conformational State of Transporter Protein OxlT

**DOI:** 10.1101/2023.10.26.564285

**Authors:** Jun Ohnuki, Titouan Jaunet-Lahary, Atsuko Yamashita, Kei-ichi Okazaki

**Affiliations:** Research Center for Computational Science, Institute for Molecular Science, National Institutes of Natural Sciences, Okazaki, 444-8585, Japan; Graduate Institute for Advanced Studies, SOKENDAI, Okazaki, Aichi 444-8585, Japan; Graduate School of Medicine, Dentistry and Pharmaceutical Sciences, Okayama University, Okayama, 700-8530, Japan

**Keywords:** Conformational transitions, Structure prediction, Biological transport, Molecular dynamics simulations, Machine learning

## Abstract

Transporter proteins change their conformation to carry their substrate across the cell membrane. The conformational dynamics are vital to understanding the transport function. We have studied the oxalate transporter (OxlT), an oxalate:formate antiporter from *Oxalobacter formigenes*, significant in avoiding kidney stone formation. The atomic structure of OxlT has been recently solved in the outward-open and occluded states. However, the inward-open conformation is still missing, hindering a complete understanding of the transporter. Here, we performed an accelerated molecular dynamics simulation to sample the extensive conformational space of OxlT and successfully obtained the inward-open conformation where cytoplasmic substrate formate binding was preferred over oxalate binding. We also identified critical interactions for the inward-open conformation. The results were complemented by the highly accurate structure prediction by AlphaFold2. Although AlphaFold2 solely predicted OxlT in the outward-open conformation, mutation of the identified critical residues made it partly predict the inward-open conformation, identifying possible state-shifting mutations.

**TOC GRAPHICS:** 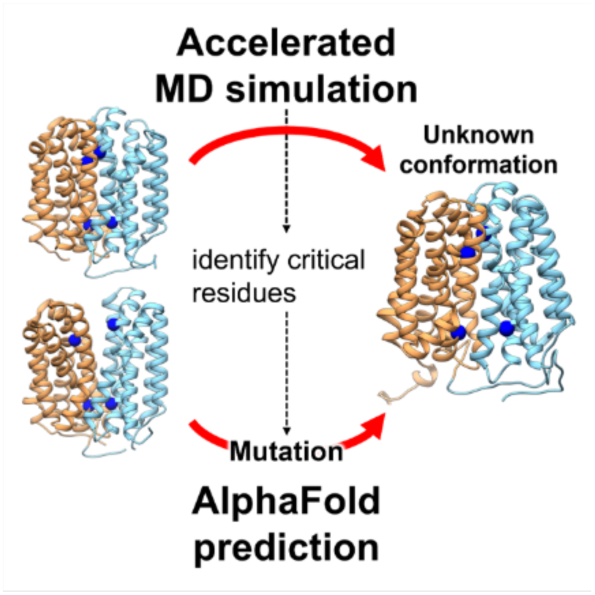

## MAIN TEXT

Oxalate (C2O4^2-^) is contained in our daily diets, such as vegetables, beans, and nuts. The oxalate level in our body is affected by the activity of an oxalate-degrading bacteria in the gut, *Oxalobacter formigenes,* that absorbs and decomposes oxalate into formate (HCO2^-^).^1^ Otherwise, excess oxalate forms an insoluble salt with calcium, which causes kidney stone disease. The oxalate transporter (OxlT), an oxalate:formate antiporter in *Oxalobacter formigenes*, is a key protein that carries oxalate inside and its product formate outside the bacterium. ^2–4^

Transporter proteins such as OxlT work by an alternating-access mechanism.^5^ They switch their conformations between the outward-open and inward-open states to alternatingly expose the binding site to opposite sides of the membrane. In the intermediate occluded conformation, access to the binding site from either side of the membrane is prohibited.^6,7^ The atomic structure of OxlT has been recently determined in the ligand-free outward-open and oxalate-bound occluded conformations (Figures 1A and 1B).^4^ However, despite structural analysis and molecular dynamics (MD) simulation, the inward-open conformation of OxlT has been missing.

**Figure 1.**
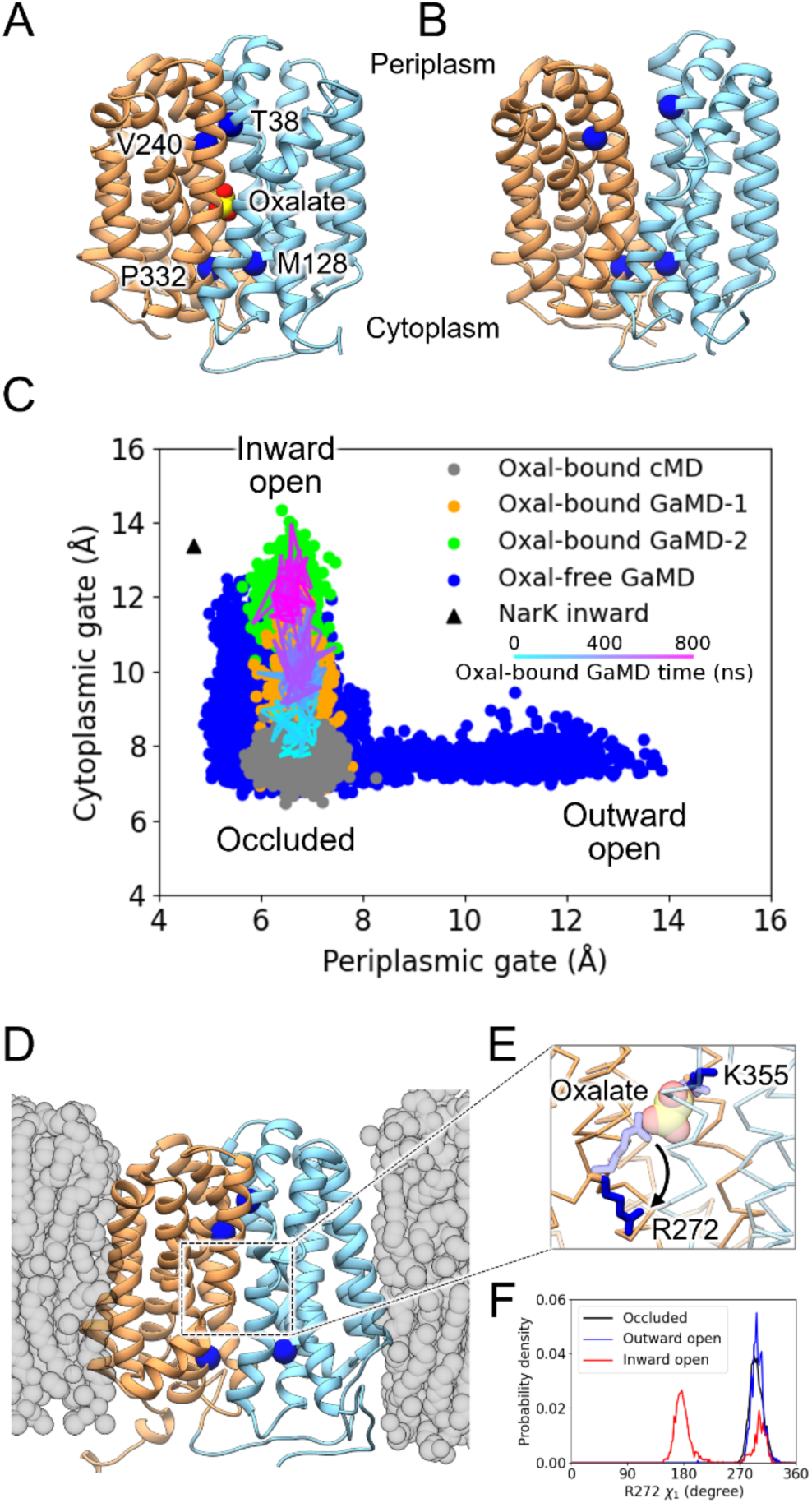
OxlT structure. (A, B) Crystal structures of oxalate-bound occluded state (A; PDB ID: 8HPK) and oxalate-free outward-open state (B; PDB ID: 8HPJ). N-terminal and C-terminal domains are colored in cyan and orange, respectively. Blue beads indicate the amino-acid residues of the periplasmic gate (T38 and V240) and those of the cytoplasmic gate (M128 and P332). (C) Distribution along the periplasmic and cytoplasmic gate distances. In oxalate-bound GaMD, boost potential was firstly applied to inter-domain interactions at the cytoplasmic region (GaMD-1) and, after 500 ns, to interactions between oxalate and the binding site (GaMD-2). Its representative trajectory is shown by a line colored from cyan (0 ns) to magenta (800 ns). The gate distances of the Nark inward open structure (PDB ID: 4U4T) are also plotted. (D) A snapshot structure of inward-open state observed in an equilibration run after the oxalate-free GaMD. The coloring is identical to (A, B). Gray beads represent the POPE lipid bilayer. (E) Magnified view of (D) to show flipping of R272 side-chain. R272, K355, and oxalate in the occluded crystal structure (transparent) are also shown for comparison. (F) Probability density distribution along χ1 dihedral of R272 in the conventional MD simulations for the oxalate-bound occluded state (black), oxalate- free inward-open state (red), and oxalate-free outward-open state (blue).

To fully understand the mechanism of OxlT, we aim to uncover the missing inward-open conformation and the crucial interaction for the conformational transition. For this purpose, we consider two computational approaches: rare-event sampling MD and highly accurate structure prediction by AlphaFold2 (AF2). On the one hand, the physics-based rare-event sampling MD can simulate slow dynamics that take longer than milliseconds, which are unreachable by conventional MD. Among several rare-event sampling methods, including the Markov state model, string method, and transition path sampling previously applied to transporter proteins,^8–10^ the accelerated MD is suited to efficiently exploring the conformational space of the transporter by boosting user-selected interaction energy.^11,12^ On the other hand, the machine-learning-based AF2 has revolutionized structural biology, achieving the highly accurate structural prediction of proteins.^13^ Although the default usage of AF2 fails to predict the conformational heterogeneity, some researchers developed a pipeline to make AF2 predict alternative conformations.^14–19^

In this study, we first sampled the extensive conformational space of OxlT by an accelerated MD simulation.^12^ After we successfully obtained an inward-open conformation, we simulated oxalate and formate binding to the inward-open OxlT, validating the conformational state, and uncovering the ligand selectivity. Then, the inter-domain contact change was analyzed between the occluded and inward-open conformation to identify critical residues for the conformational transition. The identified residues were mutated in AF2 prediction to see if they changed a bias in the conformational state.

As the initial structure for MD simulation, we employed the oxalate-bound occluded structure of OxlT (PDB ID: 8HPK).^4^ The OxlT structure was embedded into a 1-palmitoyl-2- oleoyl-phosphatidylethanolamine (POPE) lipid bilayer and solvated with water and 100 mM oxalate. All MD simulations were performed at a constant temperature of 310 K and constant pressure with anisotropic scaling using AMBER 18 or later.^20^ For the AF2-based structural prediction, the amino-acid sequence of OxlT (UniProt Q51330) was used as the wild type and modified for mutants. Multiple sequence alignment (MSA) for the sequence was generated by AlphaFold2 (ver. 2.3.1) or ColabFold^21^ (ver. 1.5.2 compatible with AlphaFold 2.3.1). Applying ColabFold to the MSA without templates, we generated 100 structure models of OxlT. Computational and methodological details are described in the Supporting Information.

To predict the unknown inward-open conformation and clarify the mechanism of the occluded-to-inward-open transition, we first performed conventional MD simulation starting from the oxalate-bound occluded state. Our previous study identified the periplasmic and cytoplasmic gates which block the water influx and the exit of oxalate in the occluded state.^4,22^ We here used the Cα-Cα distance between T38 and V240 in the periplasmic gate and the distance between M128 and P332 in the cytoplasmic gate as a measure to distinguish the conformational states of OxlT (Figure 1A). As shown in Figure 1C, the occluded conformation was stable in the conventional MD simulation, with no transition to the inward-open conformation observed.

The occluded conformation is stabilized by direct interdomain interactions as well as indirect interdomain interactions through the bound oxalate.^4^ To facilitate the transition to the inward-open conformation, we weakened the former contribution in the cytoplasmic side, i.e., the direct interdomain interaction involving the cytoplasmic gate, by using the Gaussian accelerated MD (GaMD) technique.^12^ The GaMD is an accelerated MD method that assists the system in escaping from local minima by adding harmonic boost potential to user-selected energy terms (see Computational Methods section in the Supporting Information for details).^12,23–25^ We here applied the boost potential to the cytoplasmic-side interactions between residues 128-139 of the C-terminal domain and residues 332-348 of the N-terminal domain, in which some hydrophobic and electrostatic contacts involving the cytoplasmic gate were observed.^4^ Orange points in Figure 1C show the distribution of the gate distances for 500-ns-long GaMD simulations (called “oxalate-bound GaMD-1” here). The distance of the cytoplasmic gate was increased by applying the boost potential as expected, although the distance was intermediate between the occluded conformation and the previously reported inward-open conformation of the other MFS-type transporter NarK.^26^

To clarify the role of the indirect interdomain interactions through the bound oxalate in the transition to the fully inward-open state, we then switched the boost potential from the interdomain interactions to the interactions between oxalate and the binding site (oxalate-bound GaMD-2). The cytoplasmic gate distance was increased from the intermediate distance to the fully open distance immediately after switching the boost potential (green points and the cyan-to-magenta line in Figure 1C). The result shows that the interactions both in the cytoplasmic region and in the binding site are essential to the occluded-to-inward-open transition.

In the oxalate-bound GaMD-2 simulation, although contacts of oxalate with the binding site were broken by introducing the boost potential, oxalate release to the cytoplasm was not observed in the timescale of the simulations. This is because the water channel formed in the inward-open conformation is too narrow for the bound oxalate to pass across the cytoplasmic gate. Furthermore, it cannot be ruled out that the release of oxalate to the cytoplasmic side is assisted by the entry of a formate molecule, as suggested in the exchange cycle of the NarK protein.^9^ While the binding/unbinding and translocation of oxalate is addressed later, we focus on the inward-open conformation after oxalate release (i.e., the apo state). To this end, we conducted 300-ns-long GaMD simulations for the oxalate-free occluded conformation, where the same boost potential as the oxalate-bound GaMD-1 was applied. Opening of the cytoplasmic gate was observed in multiple GaMD runs (blue points in Figure 1C), supporting the notion that the interactions of oxalate in the binding site play an important role in the transition to the inward-open conformation. We then extended the MD simulation of the oxalate-free state without the boost potential (i.e., conventional MD simulation). We verified that the observed inward-open conformation remains stable (Figures 1D and S1).

In the GaMD simulation for the oxalate-free occluded conformation (blue points in Figure 1C), we also noticed that some runs exhibited an opening of the periplasmic gate, i.e., the transition to the outward-open state. The bound oxalate stabilizes the occluded conformation by compensating for the electrostatic repulsion between the positively charged residues at the binding site (R272 and K355). Therefore, the loss of the bound oxalate destabilizes the occluded conformation and, thus, relatively stabilizes the inward-open and outward-open conformations. In the transition to the inward-open state, flipping of the R272 side chain was frequently observed (Figures 1E and 1F). This R272 flipping is likely to occur to relax the energetically unfavorable conformation due to the repulsion between R272 and K355. This is in contrast to the situation in the transition to the outward-open state where the Q34 sidechain is transiently flipped first to disrupt a hydrogen bond network at the binding site, followed by flipping of K355 to relax the electrostatic repulsion with R272.^4^ The observed difference suggests the flipping of R272 and that of Q34/K355 play a role as a direction indicator for the transition to the inward-open state and that to the outward-open state, respectively.

From the physiological function of OxlT, the ligand-free inward-open conformation is expected to bind formate from the cytoplasm, while the outward-open conformation binds oxalate from the periplasm. The inward-open and outward-open conformations obtained by the 300-ns-long oxalate-free GaMD simulation remain stable even after turning off the boost potential (Figure S1). As there is 100 mM solute oxalate in the oxalate-free OxlT system, we first investigated oxalate binding to the OxlT molecule. Figure 2A shows the spatial density distribution of water and oxalate in the inward-open conformation. No influx of oxalate into the binding site was observed in the independent six 300-ns-long MD simulation, while water filled the binding site from the cytoplasmic side. We found that oxalate was trapped at the cytoplasmic-side entrance of the water channel to the binding site, where some basic residues (R133, R139, and R284) are located. These basic residues and R272 with the side chain flipped toward the cytoplasmic side serve as an electrostatic trap to catch oxalate and prevent it from passing to the binding site (Figure 2C).

**Figure 2.**
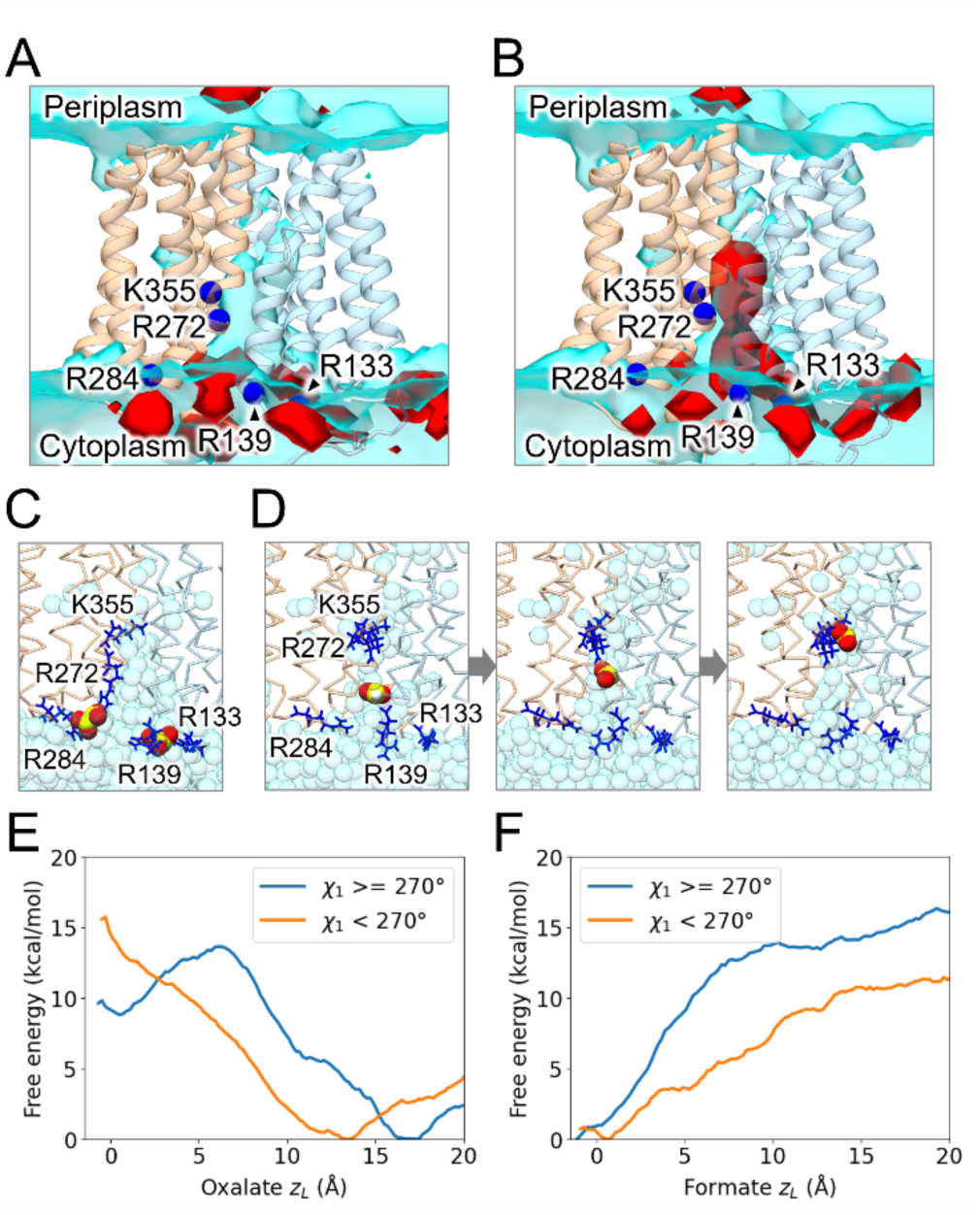
Oxalate and formate binding. (A, B) Spatial density distribution of oxalate (A) and formate (B) in the inward-open conformation. The red isodensity surface represents the relative density of oxalate or formate of 10 to the bulk. Water density is also shown in the cyan surface, corresponding to the relative density of 0.25 to the bulk water. The last 200 ns of each simulation was used for the analysis. Blue beads represent positively charges residues at the binding site and cytoplasmic entrance. (C, D) Representative snapshots of oxalate binding (C) and formate binding (D) to the inward-open conformation. Process of the formate binding is shown in three figures with a time interval of 5 ns. (E, F) Free energy profiles as a function of *zL* of oxalate (E) or formate (F) The sampling along *zL* of oxalate and formate was conducted by umbrella sampling technique, and the obtained ensemble was divided according to χ1 dihedral angle of R272.

No oxalate binding to the inward-open conformation is in contrast to the case of the outward-open conformation (Figure S2A). Our previous MD simulation showed the rapid influx of oxalate from the periplasmic side to the binding site.^4^ The periplasmic gate in the outward-open conformation can open wider than the cytoplasmic gate in the inward-open conformation (Figure S1), forming a thicker water channel than that in the inward-open conformation (Figure S2A). The asymmetry of the opening between the cytoplasmic and periplasmic gates accounts for the difference in oxalate permeability to the binding site.

We then addressed formate binding to the inward-open OxlT by replacing oxalate in water with formate. In independent six 300-ns-long MD simulations, we observed multiple binding events and a clear density distribution of formate near the binding site (Figure 2B). Formate can reach the binding site rapidly without the electrostatic trap by the basic residues at the cytoplasmic entrance (Figure 2D). This observation is explained by the less bulky and less negative charge of formate (HCO_2_^−^) than oxalate(C_2_O_4_^2−^): formate can easily escape from the trap at the cytoplasmic entrance due to weaker interactions with the basic residues and pass through the narrow water channel due to its smaller size. The filtering of substrate binding in the inward-open conformation is likely to allow *Oxalobacter formigenes* to carry formate selectively and not oxalate futilely.

The oxalate and formate binding to the inward-open OxlT was further investigated from the viewpoint of thermodynamics. To this end, we conducted an umbrella sampling^27^ to estimate the free energy profile as a function of the distance of center-of-mass of the heavy atoms of foxalate/formate from the Cα atom of K355 in the direction of the z-axis (normal to the membrane plane), denoted as *zL*. *zL* = 0 corresponds to the situation that oxalate or formate is located at the same position along the z-axis as the Cα atom of K355, and the positive direction of *zL* is pointed to the cytoplasmic side. The sampling along *zL* (0 to 20 Å) covered a wide range of the substrate position including the substrate binding site and the cytoplasmic entrance. The obtained ensemble by the umbrella sampling includes a state with R272 pointing toward the substrate binding site (*χ*1 ≥ 270°) and one with R272 pointing toward the cytoplasmic side (*χ*1 < 270°). Since the orientation of R272 side-chain is likely to affect the binding stability of oxalate/formate, we divided the ensemble according to the *χ*1 angle of R272 and obtained the free energy profile with *χ*1 ≥ 270° and with *χ*1 < 270° separately.

Figures 2E and 2F show the free energy profiles along *zL* of oxalate and along *zL* of formate, respectively. The free energy profiles clearly show that oxalate binding at the substrate binding site in the inward-open conformation is not preferred thermodynamically, regardless of the orientation of the R272 side-chain. The most stable position is found at the cytoplasmic entrance (*zL* around 17 Å for *χ*1 ≥ 270° and 13 Å for *χ*1 < 270°) of which the basic residues catch oxalate electrostatically. R272 pointing toward the cytoplasmic side (*χ*1 < 270°) is also involved in the electrostatic trap and therefore shifts the stable binding position to a more inner side of the water channel (Figure S3). In contrast, formate binding to the inward-open conformation is stable in both cases. R272 pointing toward the cytoplasmic side (*χ*1 < 270°) makes the formate binding more stable due to the interaction with formate via bridge of water molecules (Figure S3). Oxalate binding to the outward-open conformation is also preferred thermodynamically (Figure S2B). These thermodynamic properties are consistent with oxalate and formate bindings observed in the conventional MD simulations (Figures 2A, 2B, and S2A). In the case of oxalate, there is a metastable state near *zL* = 0 when R272 is pointed toward the binding site (*χ*1 ≥ 270°). This metastable state disappears upon the side-chain flipping of R272, indicating that the flipping motion of R272 promotes oxalate dissociation kinetically in the inward-open conformation.

To identify crucial interaction for the inward-open state, we analyzed changes in inter-residue contacts upon the transition from the occluded to inward-open conformation. In Figure 3A, inter-residue contacts that are strengthened (weakened) are shown in blue (red) lines. The weakened contacts are concentrated at the cytoplasmic side of the domain interface, while the strengthened contacts are at the periplasmic side of the domain interface. Especially, D78, R272, and D280 are central residues of the weakened contacts upon the transition to the inward-open conformation. The two aspartate residues D78 and D280 form charge-dipole interactions with the N-terminal ends of TM11 and TM5, respectively, as well as ionic interactions with positively charged residues at the cytoplasmic side in the occluded conformation (Figure 3B),^4^ which are lost upon the transition to the inward-open state (Figure 3C). The breaking of the interactions formed by D78 and D280, part of the motif A conserved in MSF proteins,^4^ is crucial for the transition to the inward-open conformation. It is also worth noting that our MD simulations and contact analysis captured a subtle but significant strengthening of inter-domain contacts on the opposite side of the opened cytoplasmic gate (i.e., periplasmic side). Although changes in the opposite side have not been reported in previous studies on MFS proteins, tighter packing of the periplasmic side in the inward-open conformation is plausible for OxlT to work rigorously by the alternating-access mechanism.

**Figure 3.**
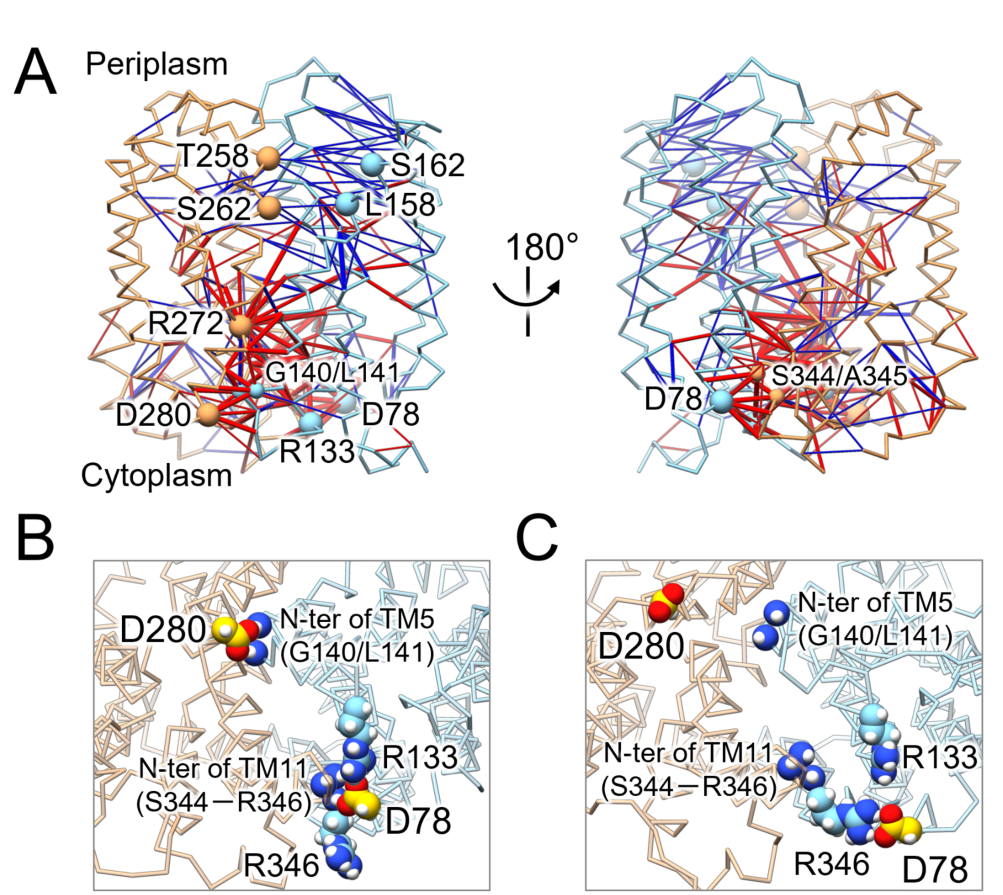
(A) Inter-residue contact changes upon the transition to the inward-open conformation. Blue and red bonds represent the contacts of which probability increases and decreases, respectively. A residue pair was considered in contact if the closest distance between any two heavy atoms in the residues was less than 4.5 Å. The contact probability for the occluded conformation, *p*_o_, was calculated from snapshot structures after 100 ns of the five 1-µs-long conventional MD simulations of the oxalate-bound occluded conformation. The contact probability for the inward-open conformation, *p*_i_, was calculated from snapshot structures after 100 ns of the six 300-ns-long conventional MD simulations of the oxalate-free inward-open conformation. The bond thickness reflects the magnitude of the probability change (|*p*_i_ − *p*_o_|). We omitted the bonds with |*p*_i_ − *p*_o_| lower than 0.2 or (δ*p*_i_^2^ + δ*p*_o_^2^)^1/2^, where δ*p*_i_ and δ*p*_o_ are the standard errors of *p*_i_ and *p*_o_, respectively. Domain coloring is the same as Figure 1. (B, C) View from the periplasmic side showing snapshots of the occluded (B) and inward-open (C) conformations.

The identified crucial interactions for the inward-open state are now complemented by the highly accurate structure prediction by AF2.^13^ Because AF2 only learns and predicts the final structures of protein folding ^32^, there is difficulty in predicting the conformational heterogeneity. The prediction is usually biased to one conformation among possible conformations taken during the functional cycle,^33,34^ which has been recently addressed by MSA subsampling,^15^ masking,^16^ or clustering^17^ and template-based prediction.^18,19^ We first generated an MSA from the wild-type sequence of OxlT with JackHMMer^28^ and HHblits^29^ used in AF2. Applying AF2 with the MSA, we found that all predicted models are in the outward-open conformation (black dots in Figure 4A), which coincides with the previous structural determination that found OxlT in the outward-open conformation in the absence of ligand.^4^ Note that the OxlT outward-open structure was not included in the database of the AF2 prediction.

**Figure 4.**
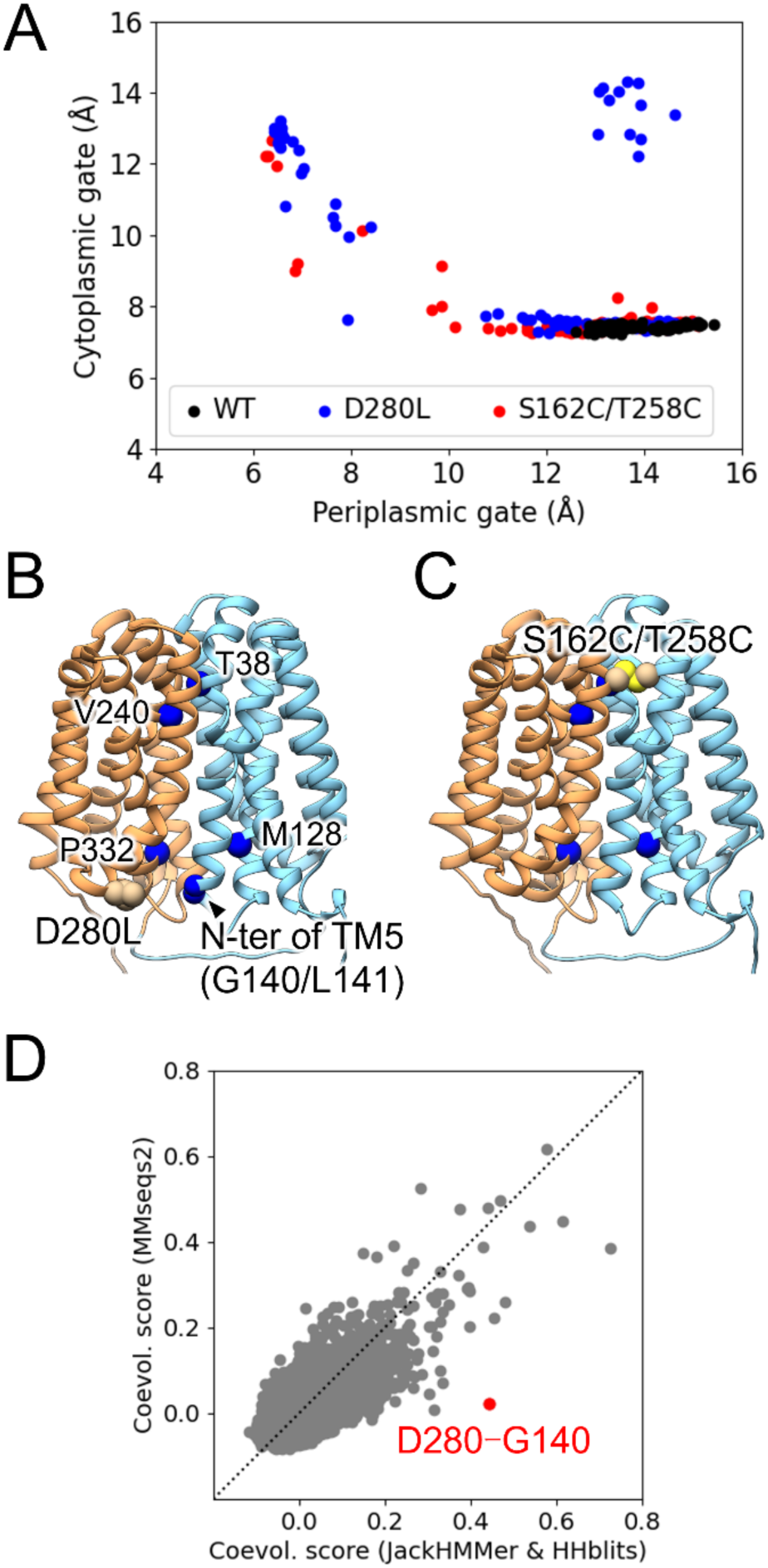
AF2 structure prediction. (A) Periplasmic and cytoplasmic gate distances of wild-type and mutant OxlT structures by AF2. (B) Structure of D280S mutant. (C) Structure of T258C/S162C double mutant. (D) Coevolution score for MSA with JackHMMer^28^ and HHblits^29^, versus MSA with MMseqs2.^30^ Coevolution score was obtained by pseudo-likelihood maximization direct coupling analysis.^31^ A higher score for an amino-acid residue pair indicates a stronger coevolution coupling.

Then, we sought mutations that can shift the bias to the inward-open conformation. Based on the inter-domain contact analysis of MD simulation (Figure 3), we focused on D280 in the A motif at the cytoplasmic side and S162/T258 at the periplasmic side. The inter-domain contacts of D280 with the N-terminal of TM5 (G140/L141) were lost upon the transition from the occluded to inward-open conformation (Figures 3B and C). The D280L mutation was intended to diminish the contacts formed in the occluded conformation, thus shifting toward the inward-open conformation. For S162/T258, the distance of the residue pair becomes closer upon the transition. The double mutation S162C/T258C was intended to form a disulfide bond between the two in the inward-open conformation. For both D280L and S162C/T258C mutations, AF2 partially predicted the inward-open conformation, shown in blue and red points in Figure 4A, respectively. As shown in the AF2 structures (Figures 4B and C) and gate distances (Figure 4A), these mutations remarkably showed a conformational shift to the inward-open conformation. The D280L mutation, however, occasionally leads to a loose conformation with both periplasmic and cytoplasmic gates open. The state-shifting mutations should be helpful for experimental structural determination of the missing conformational state.

We further examined why the bias to the outward-open conformation was observed for the wild-type OxlT. The previous studies showed that MSA used in AF2 affects the structure prediction.^15–17^ Here, we generated another MSA with MMseqs2^30^, which is used in ColabFold^21^. Surprisingly, ColabFold with the MSA predicted the conformational diversity, including the outward-open, occluded, and inward-open conformations (Figure S4A). To figure out the reason for the different results, we applied the direct coupling analysis^31^ to the two MSAs. The coevolution analysis showed a higher coevolution score for cytoplasmic inter-domain residue pairs in the MSA with JackHMMer^28^ and HHblits^29^ (Figure S4B) compared to those in the MSA with MMseqs2^30^, which can be seen in the lower-right part of Figure 4D. D280-G140 is the most prominent among these pairs. This result suggests that AF2 has a higher weight for the cytoplasmic inter-domain contacts, which leads to the biased prediction of the outward-open conformation. The result also explains why the D280 mutation shifts toward the inward-open conformation. By disrupting the coevolved cytoplasmic inter-domain contacts, the mutation made AF2 partly predict the inward-open conformation. The recent coevolution analysis of sugar porters by Drew, Delemotte, and coworkers also showed the importance of state-dependent coevolving residue pairs.^35^

In conclusion, we uncovered the inward-open conformation of OxlT by the accelerated MD simulation, filling the missing piece in the OxlT transport cycle. The obtained inward-open conformation was then subjected to the oxalate/formate binding simulations. The simulations showed selective binding of formate to the inward-open conformation, while oxalate was blocked at the entrance of the binding site. This result provides the physical basis for the oxalate/formate antiport function of OxlT: After oxalate is transported to the cytoplasmic side of the membrane, formate, but not oxalate, binds to the inward-open OxlT and is transported to the periplasmic side of the membrane. Finally, we integrated the physics-based MD results with the machine-learning-based AF2 structure prediction. The AF2 prediction of the wild-type sequence all resulted in the outward-open conformation of OxlT. We introduced single/double mutations to the residues involved in crucial interactions of the inward-open state. Among these mutations, the D280L mutation was intended to disrupt the contacts formed in the occluded conformation at the cytoplasmic side, and the S162C/T258C mutation was intended to form a disulfide bond at the periplasmic side in the inward-open conformation. We found that the mutations shift OxlT to the inward-open state. Although previous studies suggested that AF2 is unable to predict point mutations that cause misfolding or destabilization of the native structure,^36,37^ the situation is different for the protein conformational change because AF2 can learn alternative conformations of homologous proteins in PDB. A previous study also showed a possibility that two or more mutations allow AF2 to explore an alternative conformation of transporter proteins.^38^ In combination with the accelerated MD simulation, we here clarified that even a single mutation can shift AF2-predicted conformation. Furthermore, we showed that AF2 prediction is affected by the input MSA, and residue-wise coevolution score derived from the MSA can pin down the state-shifting mutations. Our general procedure can be applied to other proteins, predicting conformational-state-shifting mutations.

## ASSOCIATED CONTENT

## AUTHOR INFORMATION

Notes

The authors declare no competing financial interests.

## ACKNOWLEDGMENT

The computation was partially performed using Research Center for Computational Science, Okazaki, Japan (Project: 22-IMS-C189 and 23-IMS-C201). This work was supported by JSPS KAKENHI Grant Numbers JP23K14160 (to J.O.), JP22H02595 (to K.O.) and NAGASE Science and Technology Foundation (to A.Y.).

