## Supporting Information for "Accelerated Molecular Dynamics and AlphaFold Uncover a Missing Conformational State of Transporter Protein OxlT"

AUTHOR INFORMATION

**Corresponding Author**

*

### Computational Methods

*System preparation*

We employed the oxalate-bound occluded structure of OxlT (PDB ID: 8HPK)[1] as the initial structure for MD simulation. Molecules included in the structure were eliminated except for OxlT and the bound oxalate. Modeller[2] was used to model missing residues at the N-terminal tail, the central loop, and the C-terminal tail except for the initiator methionine. The OxlT structure was embedded into a 1-palmitoyl-2-oleoyl-phosphatidylethanolamine (POPE) lipid bilayer in a box with the initial size of 120×120×105 $Å^3$, where the lipid bilayer was aligned parallel to the x－y plane, and solvated with water and 100mM $K_2CO_3$ by using the Membrane Builder plugin in CHARMM-GUI.[3,4] All $CO_3^{2-}$ were replaced with oxalate ions. According to the calculated p*K*a by H++,[5] His418 was neutralized, and the other basic residues and the acidic residues were charged. We used the ff14SB force field for OxlT,[6] the lipid14 force field for POPE,[7] the TIP3P water model,[8] and the potassium parameter by Joung and Cheatham.[9] For the bound and solvent oxalates, we used the parameter prepared for the bound oxalate in our previous study,[1] where bonding parameters were based on the Ab Initio MD simulation by Kroutil et al.[10,11] and partial charges were determined by RESP (Restrained Electrostatic Potential) scheme.[12]

*Conventional MD simulation of occluded OxlT in oxalate-bound state*

The periodic boundary condition was applied to the solvated box. Long-range electrostatic interactions were calculated by particle mesh Ewald method[13] with a direct space cutoff of 10 Å. The time step was set to 2 fs by applying the SHAKE method[14] to bonds involving hydrogen. The simulation conditions described in this paragraph are also used in the following all MD simulations including Gaussian accelerated MD and umbrella sampling.

The oxalate-bound OxlT system was energy minimized (1000-step steepest descent and subsequent 1000-step conjugate gradient minimizations) first in the presence of positional restraints applied to OxlT and the bound oxalate (force constant of 500 kcal/mol/Å$^2$) and second in the absence of the restraints. Then, the system was heated up to 100 K at constant volume for 5 ps where the Langevin thermostat[15] was used with a collision frequency of 1 ps$^{-1}$. At this stage, we applied the positional restraints to OxlT, the bound oxalate, and the lipid (force constant of 10 kcal/mol/Å$^2$). Then the temperature was increased to 310 K and the volume was relaxed for 100 ps at 1 bar without restraints where the pressure was controlled anisotropically by the Monte Carlo barostat.[16,17] Finally, we conducted five independent 1-µs-long product runs at 310 K and 1 bar.

*Gaussian accelerated MD simulation*

To enhance the transition from the occluded conformation to the inward-open conformation, we selected two snapshot structures at 500 ns from the five conventional MD runs and conducted the Gaussian accelerated MD (GaMD) simulation.[18] GaMD and its variants[19–21] adds the following boost potential to the selected term of the system potential energy:

$$\Delta V(\boldsymbol{r}) = \begin{cases} 0 & (V(\boldsymbol{r}) \geq E) \\ \frac{1}{2}k\left(E - V(\boldsymbol{r})\right)^2 & (V(\boldsymbol{r}) < E) \end{cases} \quad (1)$$

where $V(r)$ is the user-selected energy term, $k$ is the force constant, and $E$ is the threshold energy. Thus, the boosted energy $V^*(r)$ is given by

$$V^*(r) = V(r) + \Delta V(r) \quad (2)$$

As the energy term to which the boost potential is applied, we here selected interactions between residues 128-139 of the C-terminal domain and residues 332-348 of the N-terminal domain that correspond to the inter-domain interface around the cytoplasmic gate. In order to ensure the smoothed energy surface and the efficient sampling, we determined the parameters $k$ and $E$ as satisfying the criteria suggested by Miao and colleagues[18]:

$$k = \frac{k_0}{(V_{max} - V_{min})}$$

$$0 < k_0 \leq 1$$

$$k_0 \geq \left(1 - \frac{\sigma_0}{\sigma}\right) \frac{V_{max} - V_{min}}{V_{avg} - V_{min}}$$

$$E = V_{min} + \frac{1}{k} \qquad (3)$$

where $V_{min}$, $V_{max}$, $V_{avg}$, and $\sigma$ are the minimum, maximum, average, and standard deviation of the selected energy term. The parameter $\sigma_0$ in the criteria was set to 6.0 kcal/mol. The energy statistics ($V_{min}$, $V_{max}$, $V_{avg}$, and $\sigma$) and the boost parameters ($k$, $E$) were estimated from the short MD simulations: (i) 2-ns-long conventional MD where the statistics were calculated from the energies collected after 0.4 ns, (ii) 2-ns-long GaMD where the boost parameters were firstly determined from the statistics at stage (i) and then updated from the energies collected after 0.4 ns. Because the right-hand side in the third equation of eq. (3) was lower than zero, we set $k_0$ = 1.0 and accordingly $E = V_{max}$. By using the parameters, we conducted two independent 500-ns-long GaMD simulations. In addition, to investigate the role of the bound oxalate in the transition to the inward-open conformation, we further extended one GaMD run to 800 ns by switching the boosted energy term at 500 ns from the inter-domain interactions to interactions between oxalate and the binding site (Q34, Y35, Y124, A147, R272, Y328, W352, and K355). We also prepared the oxalate-free OxlT system by removing the bound oxalate from the 500 ns snapshot structures of the oxalate-bound system and 25 independent 300-ns-long GaMD simulations. All GaMD simulations were carried out at 310 K and 1 bar where we used the Langevin thermostat[15] with the collision frequency of 1 $ps^{-1}$ and the anisotropic Berendsen barostat[22] with the pressure relaxation time of 5 ps.

*Equilibration of inward-open or outward-open OxlT in oxalate-free state*

As mentioned in the main text, the GaMD simulation of the oxalate-free OxlT captured the transition to the inward-open conformation as well as that to the outward-open conformation. By conducting additional conventional MD simulations, we investigated if the obtained inward-open and outward-open conformations remain stable without the boost potential and if solvent oxalate molecules enter into and bind with the binding site of OxlT. As the starting structures, we selected the final structures of the oxalate-free GaMD simulations where the transition to the inward or outward-open conformation was observed. In addition, we also prepared the system with formate by replacing oxalate with formate, where the number of potassium ions was adjusted to ensure charge neutrality. For the system with oxalate, we conducted six 300-ns-long MD simulations for the inward-open conformation and three 300-ns-long MD simulations for the outward-open conformation. For the system with formate, we conducted six 300-ns-long MD simulations for the inward-open conformation.

*Free energy calculation of oxalate and formate position*

To clarify the oxalate and formate transport by the inward-open OxlT from the viewpoint of thermodynamics, we conducted the umbrella sampling[23] for $z$ component of the center-of-mass distance of ligand (i.e., oxalate or formate) from the $C_\alpha$ position of K355, denoted by $z_L$. The $z$ axis is perpendicular to the lipid membrane plane. The following umbrella potential was applied to $z_L$,

$$U(z_L) = K(z_L - z_0)^2 \quad (4)$$

where $z_0$ is the equilibrium position and $K$ is the force constant. $z_0$ was shifted from 0 to 20 Å at the intervals of 0.5 Å (41 windows). $K$ was set to 2.5 kcal/mol/Å$^2$. As the initial system, we employed the final structures of the conventional MD simulations in the oxalate-free state. One oxalate (formate) molecule in proximity to OxlT was used for sampling along $z_L$, and the others were replaced with chloride ions, where the number of potassium ions was adjusted to ensure charge neutrality. To prepare the starting structure for each window, the selected oxalate (formate) was moved from its initial position toward $z_L$ = 0 Å or toward $z_L$ = 20 Å by the harmonic potential with the same form as eq. (4) where $z_0$ is the time-dependent equilibrium position changing at a speed of 1 Å/ns and $K$ is the force constant set to 3.1 kcal/mol/Å$^2$. Finally, we conducted six 16-ns-long MD simulations for each window of the inward-open conformation and three 16-ns-long MD simulations for each window of the outward-open conformation. Free energy profile along $z_L$ was obtained from reweighting of the probability density c by the weighted histogram analysis method (WHAM).[24,25]

*Structural prediction by AlphaFold2 and ColabFold*

AlphaFold2[26] (ver. 2.3.1) and ColabFold[27] (ver. 1.5.2 compatible with AlphaFold 2.3.1) were used for structural prediction of OxlT. The amino-acid sequence of OxlT (UniProt Q51330 residues 2－418) was used as the wild type and modified for the D280S and S162C/T258C mutants. Multiple sequence alignment (MSA) for the sequence was generated by JackHMMer[28] and HHBlits[29] through AlphaFold2 and was provided for ColabFold to generate 100 structure models (20 random seeds were used where each seed generated five models). Templates were not used in the structural prediction by ColabFold. For the depth of MSA subsampling, we set the number of sequence clusters 'max_msa_clusters' to 512 and the number of extra sequences 'max_extra_msa' to 1024. The number of prediction recycles was set to 3, and structural relaxation after prediction was not performed. We also performed ColabFold both to generate MSA and to predict the OxlT structure in which MSA generation was executed using MMseqs2[30] rather than JackHMMer and HHBlits and investigated the effect of MSA on the structural prediction.

Inter-residue coevolution coupling in MSA was investigated by pseudo-likelihood maximization direct coupling analysis[31] using the software "pydca".[32] To calculate the coevolution score, we used the Frobenius norm of the inter-residue couplings and average product correction.

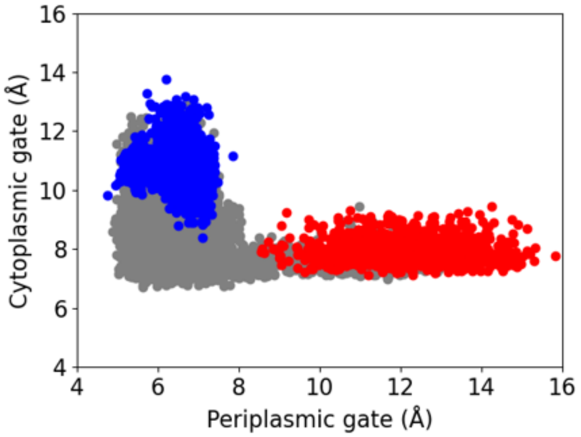


**Figure S1.** Periplasmic and cytoplasmic gate distances in oxalate-free conventional MD (cMD) simulations. As mentioned in the main text, oxalate-free GaMD simulation captured transition events toward the inward-open conformation or toward the outward-open conformation (gray). Starting from the obtained inward-open and outward-open conformations, we extended the oxalate-free MD simulation without boost potentials, i.e., cMD simulation (blue: cMD starting from the inward-open conformation, red: cMD starting from the outward-open conformation).

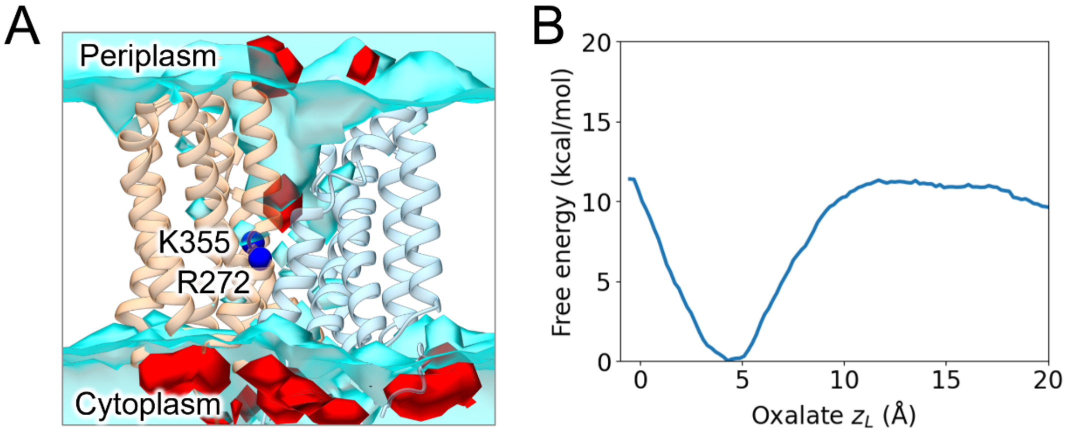


**Figure S2.** (A) Spatial density distribution of oxalate in the outward-open conformation. The red isodensity surface represents the relative oxalate density of 10 to the bulk. Water density distribution is also shown in the cyan isodensity surface, corresponding to the relative density of 0.25 to the bulk water. (B) Free energy profile as a function of $z_L$ ($z$ component of the distance of oxalate from the $C_\alpha$ position of K355) in the outward-open conformation.

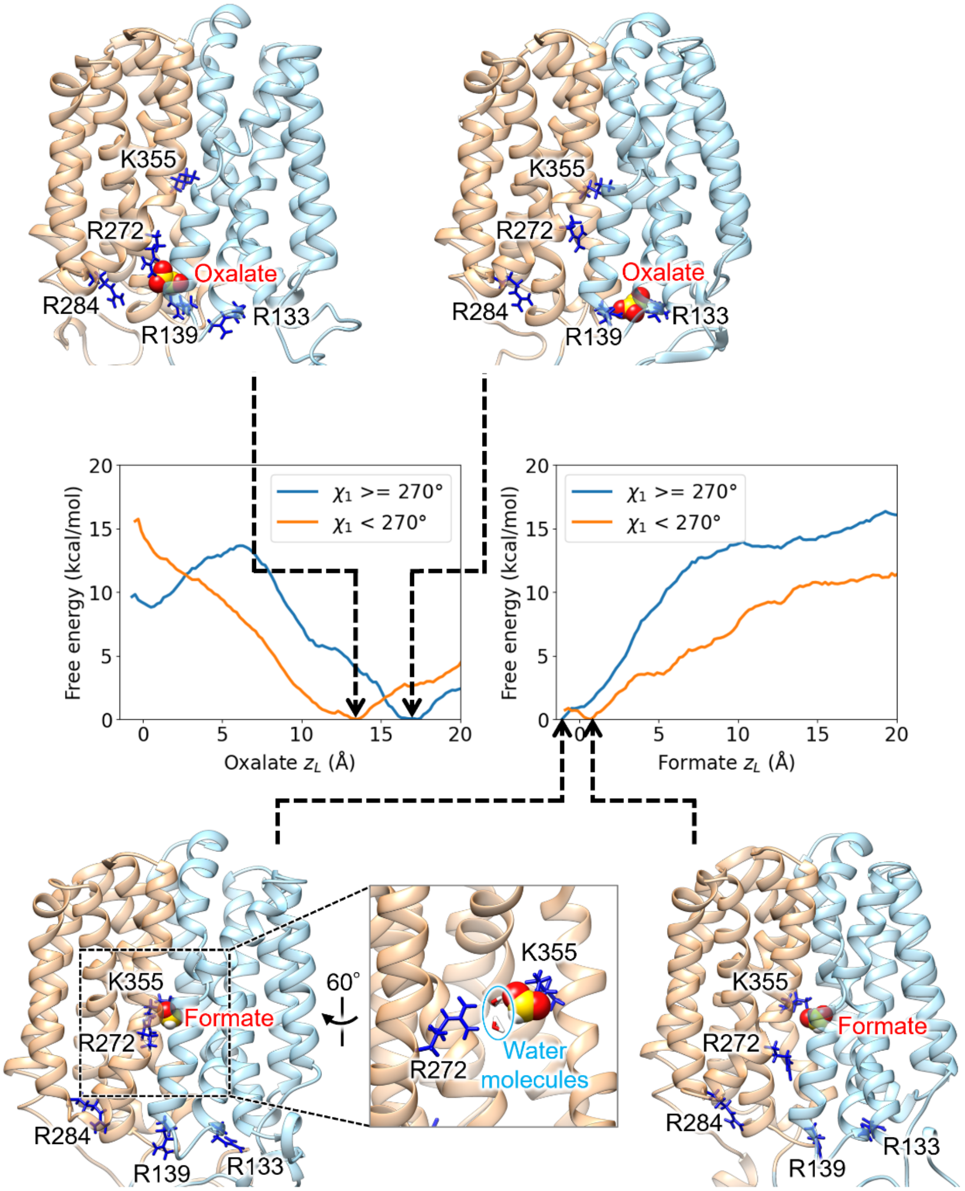


**Figure S3.** Binding mode of substrate at the most stable region of free energy profile.

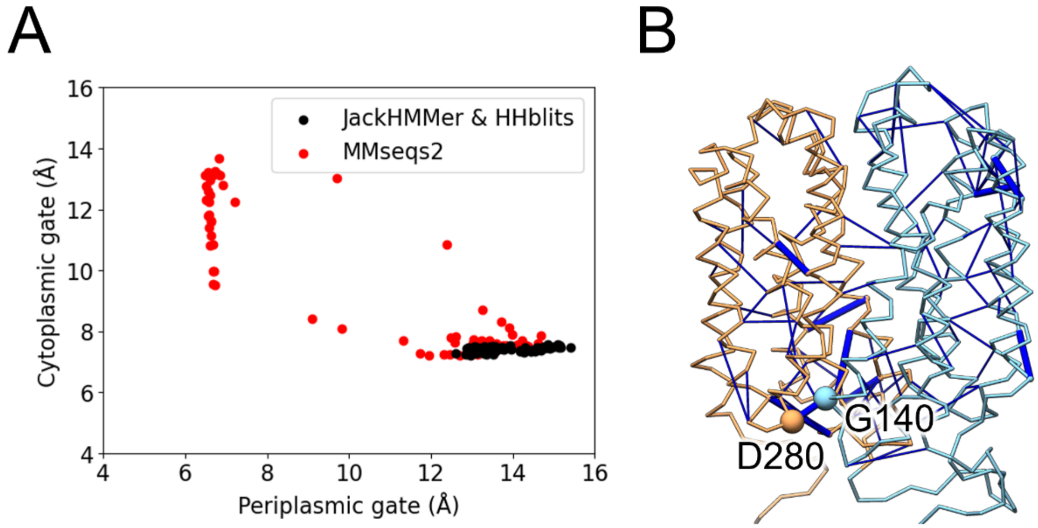


**Figure S4.** (A) Periplasmic and cytoplasmic gate distances of the OxlT structures predicted from MSA with JackHMMer[28] and HHblits[29] (black) and from MSA with MMseqs2[30] (red). (B) Residue pairs with top 0.01% (thick blue sticks) or 0.1% (thin blue sticks) coevolution score. Coevolution score was obtained applying the direct coupling analysis to the MSA with JackHMMer and HHblits.
